# SpeciesMLP: Sequence based Multi-layer Perceptron For Amplicon Read Classification Using Real-time Data Augmentation

**DOI:** 10.1101/419846

**Authors:** Ali Kishk, Mohamed El-Hadidi

## Abstract

Taxonomic assignment is the core of targeted metagenomics approaches that aims to assign sequencing reads to their corresponding taxonomy. Sequence similarity searching and machine learning (ML) are two commonly used approaches for taxonomic assignment based on the 16S rRNA. Similarity based approaches require high computation resources, while ML approaches don’t need these resources in prediction. The majority of these ML approaches depend on k-mer frequency rather than direct sequence, which leads to low accuracy on short reads as k-mer frequency doesn’t consider k-mer position. Moreover training ML taxonomic classifiers depend on a specific read length which may reduce the prediction performance by decreasing read length. In this study, we built a neural network classifier for 16S rRNA reads based on SILVA database (version 132). Modeling was performed on direct sequences using Convolutional neural network (CNN) and other neural network architectures such as Multi-layer Perceptron and Recurrent Neural Network. In order to reduce modeling time of the direct sequences, In-silico PCR was applied on SILVA database. Total number of 14 subset databases were generated by universal primers for each single or paired high variable region (HVR). Moreover, in this study, we illustrate the results for the V2 database model on 8443 classes on the species level and 1552 on the genus level. In order to simulate sequencing fragmentation, we trained variable length subsequences from 50 bases till the full length of the HVR that are randomly changing in each training iteration. Simple MLP model with global max pooling gives 0.71 & 0.93 test accuracy for the species and genus levels respectively (for reads of 100 base sub-sequences) and 0.75 & 0.96 accuracy for the species and genus levels respectively (on the full length V2 HVR). In this study, we present a novel method (SpeciesMLP https://github.com/ali-kishk/SpeciesMLP) to model the direct amplicon sequence using MLP over a sequence of k-mers faster 20 times than CNN in training and 10 times in prediction.

**Figure 1:**
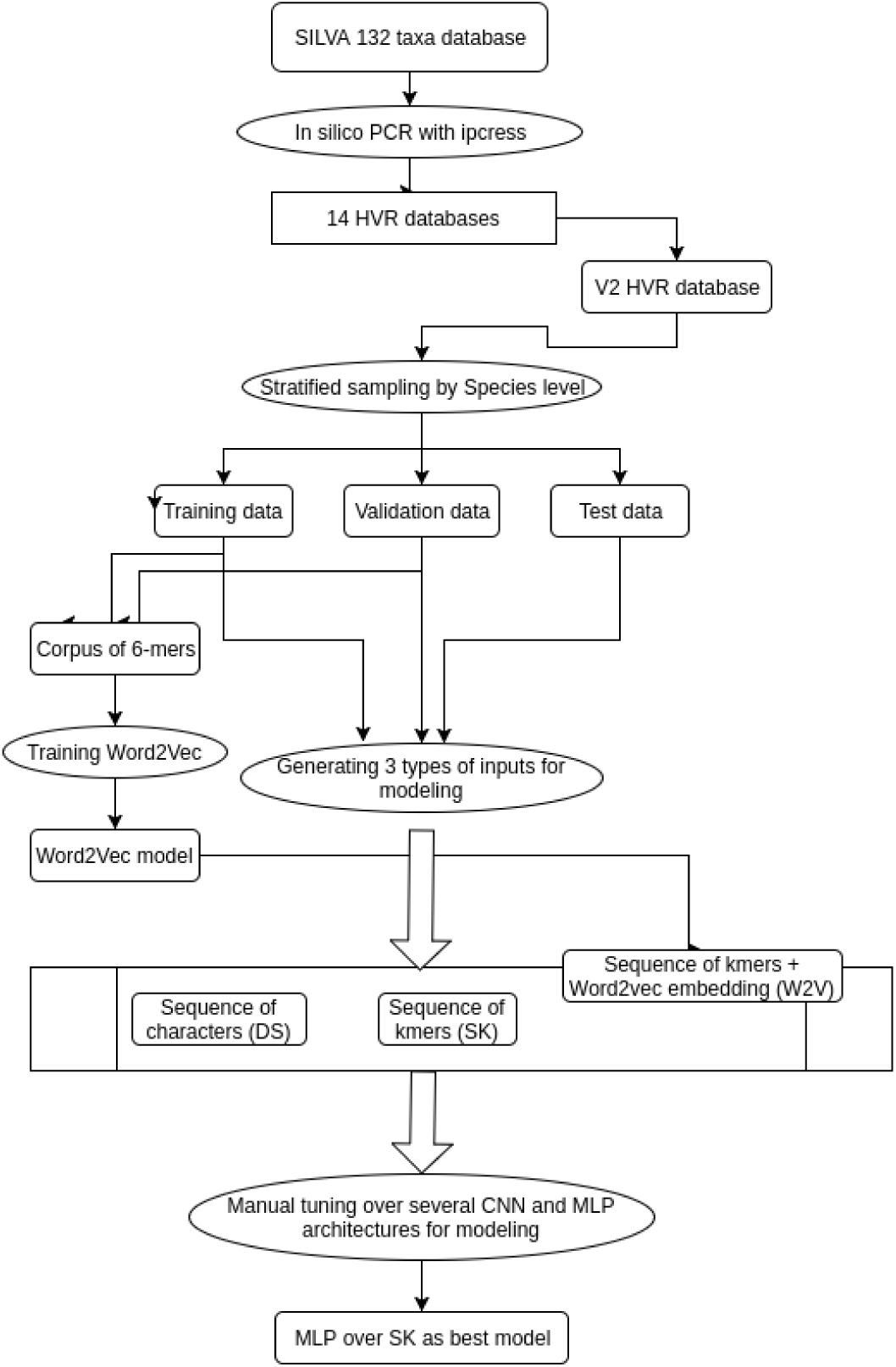
Workflow diagram. The diagram begins with Insilico PCR to SILVA taxa. The rest pipeline is applied only to the V2 HVR database only from splitting to modeling.

## I. Background

Metagenomics is the study of a microbial communities’ genetic fingerprint. Recently it was shown that these communities contribute to many biological functions such as antibiotic resistance [1], metabolism [2] and immunity regulation [3]. Regarding its role in the environment, they have important roles in climate change [4], the space environment [5], water biofilm [6] and nuclear waste detoxification [7]. As metagenomics is changing our insight on health and the environment, new sequence analysis methods are becoming more important.

Metagenomics data are generated from two main sequencing techniques, targeted sequencing [8], and whole genome shotgun sequencing [8]. Targeted or amplicon sequencing depends on PCR amplification of the rRNA genes especially 16S rRNA for bacteria and archaea, while shotgun sequencing uses the sequence of the whole genome. Amplicon sequencing allows cheap and rapid identifications of the microbial species whereas shotgun can define the new microbial content and allow another dimension for functional classification [8].

Machine learning is the ability of computers to learn from the data without any stored instructions. ML algorithms are mainly two types: supervised and unsupervised. Supervised ML is modeling data with known outputs feature for each sample where unsupervised ML don’t. Supervised ML tries to map an output to a set of inputs, where regression is a supervised ML in case the input is numerical, whereas classification in case of categorical input. Deep learning (DL) is a subtype of ML in both supervised and unsupervised, where it utilizes the connectivity and architecture of the human nervous system for modeling. The revival of DL allowed modeling of complex natural data such images, audio, and text, where it was hard to model using classical ML algorithms such as Random Forest [9] and Support Vector Machine [10].

## II. Introduction

### A. Metagenomics analysis steps

Both whole genome shotgun and amplicon sequencing methods share some bioinformatics algorithms in analysis such as quality processing and taxonomic assignment. Taxonomic assignment [11] is usually the final step of metagenomic analysis and probably the most computationally demanding. It involves assigning each read in each sample to a corresponding taxonomy on many levels from phylum to species or even strain level.

### B. Taxonomic assignment

Taxonomic assignment uses alignment or machine learning for classification. Alignment based approaches such as BLAST [12], MALT [13], and SINA [14] need a reference 16S rRNA database for sequence comparison. ML-based approaches such as RDP [15], 16S Classifier [16] need first to be trained on the reference database to produce a classifier. Once trained, these classifier models can be used for taxonomic assignment. Alignment based approaches are generally more accurate than ML-based but at the expense of the computational resources [17].

### C. Deep Learning in Bioinformatics

DL application covers many bioinformatics fields including genomic [18], transcriptomics [19]. metagenomics [20] and other OMICs fields. Data used in these applications can be classified into a feature-based or direct sequence based.Phenotype classification such as classifying samples to healthy and diseased is the most widespread application of feature-based DL. Phenotype classification can be found in genomics from mutation data [18], transcriptomics from differentially expressed genes [19] and metagenomics from Operating Taxonomy Unit (OTU) tables [21] or k-mer frequency [20]. The ability of DL to model thousands to millions of features allowed direct modeling without feature selection. Multi-layer perceptron (MLP) [22] is the DL architecture commonly used in phenotype classification [20].

### D. Deep Learning in Metagenomics

Phenotype classification and gene ontology prediction [23] are among commonly used DL applications in metagenomics. All phenotype classification methods are feature-based such as MicroPheno depends on k-mer frequency with MLP. Others depend on OTUs tables instead of k-mer frequency such as [21] used MLP and recursive neural network for phenotype classification and learning the hierarchy between the samples. Also, [24] used CNN [25] to learn the hierarchy between the OTUs and phenotype classification. Moreover, [26] used CNN on images generated by phylogenetic sorting on the OTUs for each sample for phenotype classification. DeepARG [23] is an example of gene ontology application using MLP for detecting antibiotic resistant genes.

### E. Word2Vec in Bioinformatics

Some DL techniques allow faster sequence classification such as Word2Vec [27]; a simple MLP hidden layer where the inputs are the context k-mers and the output is a target k-mer. Word2Vec is a self-supervised learning method that allows close target k-mers from the concept of replaceability in a specific context of k-mers to have close vectors. Empirically Word2Vec enhances any supervised learning such as classification [28].

## III. Related Work

RDP [15] is the first ML taxonomic classifier using Naive Bayes classification on k-mer frequency. RDP was trained on 300K sequences from Bergey’s Taxonomic Outline [29] with a k-mer size of 8 from phylum to genus level. 16S Classifier [16] applied Random Forest for classification on Greengenes [30], where In-silico PCR was used to generate HVR subset databases to reduce the search space. All possible k-mers with size ranges from 2 to 6 were used, then feature selection was applied to reduce them. 16S Classifier produced specific models for each HVR database.

As new DL architectures develop rapidly such as CNN and Recurrent Neural Network (RNN) [31], direct sequence based DL approaches are dominating over feature based approaches. CNN was developed mainly for visual computing but used later in sequence classification. CNN finds temporal features where RNN searches for consecutive features in a sequence. RNA [32] and protein [33] structure prediction and alternative splicing prediction are among direct sequence based applications. The only available example (by mid-June 2018 besides this study) of sequence-based DL application in metagenomics is [34] for OTUs clustering using CNN and PCA.

Word2Vec was recently applied in alternative splicing prediction in SpliceVec [28], where it replaced complex DL architectures with MLP and to increase the model accuracy. Additionally, Word2Vec has used in medical text mining [35] and protein structure classification [36].

## IV. Materials & Methods

### A. Generating HVRs databases

In order to decrease the search space by the classifier model, In silico PCR was applied on SILVA database [37] (version 132, SILVA_taxa, accessed in April 2018) to generate a subset for each HVR (eg: V2) or pairs of HVRs (eg: V3-V4). Pairs of primers for each HVR subset were the same in the 16S Classifier [16], all ambiguity characters in any primer sequences were expanded using an in-house developed script. The expended primer database was used in the In silico PCR by ipcress (version 2.2.0) [38] against each HVR database. All next steps were applied separately for each HVR dataset.

### B. Taxonomy parsing

Taxonomy ranking from phylum to species was parsed for each sequence. We discard any sequence with at least one missing rank (eg: order is missing in the hierarchy). In some cases, there are reads with unknown genus or species among different families, in order to avoid confusion, their previous ranks were concatenated to their genus and species classes.

### C. Ambiguity character handling

As SILVA database contains ambiguity characters rather than A, C, T, and G, any sequence with ambiguity characters was expanded to only one example. Expanding ambiguity sequences by more than one example resulted later in class in-balance and fast over-fitting. Biopython (version 1.72) [39] was used in taxonomy parsing and Ambiguity character handling.

### D. Stratified sampling

Training, test and validation datasets were split in a stratified manner using the species level as the output. Further preprocessing was done to ensure the same species classes exist in the 3 datasets.

### E. Word2Vec model training

As Word2Vec model deals with words rather than characters, each sequence of nucleotide characters was converted to a sequence of k-mers with k-mer size=6. A corpus of k-mers was generated from the validation and training data. Word2Vec was trained for 5 epochs using the skip-gram algorithm from the locally saved corpus. Gensim package (Version 3.4) [40] was used in training the Word2Vec model.

### F. Classifier training

#### a) Architectures

Several DL architectures in text classification were tested such as CNN, RNNs & MLP. Among tested CNN architectures are Residual network (ResNet) [41] & Inception [42]. The tested CNN architectures use 1D convolution instead of 2D convolution. RNNs are the least explored models such as Gated Recurrent Unit [43] and Long Short-Term Memory [44]. All models end with 6 fully connected layers, each corresponding to each rank from phylum to species. Species-level output has the higher number of classes. Weighted Adam optimization [45] was applied with early stopping if the model didn’t coverage within 5 epochs. Manual architecture tuning was done on V2 HVR only as an example for the rest HVR databases. All the networks were built using Keras (Version 2.2.0) [46].

#### b) Real-time variable length modeling

In order to simulate variability in read lengths generated by NGS, a custom training function generated variable length reads was applied. This function generated sub-sequences of the original ones that are changing in each training epoch. The maximum length was 320 as it’s the maximum read length in V2 HVR dataset. The minimum length was set to 50. To compare the effect of variable length against fixed length data augmentation, we applied the same hyper-parameters of our best model for training a fixed length reads of length 100 bases.

#### c) Input representation & Model Evaluation

For the Word2Vec, 3 types of input were tested: the direct sequence of characters (DC), the sequence of k-mers with Word2Vec embedding (W2V) and the sequence of k-mers without Word2Vec (SK). A range of read lengths (25, 50, 75, 100, 125, full_length HVR) was used to simulate fixed-length reads data on the test data for each length. Training time and the maximum test accuracy are the parameters in choosing of the best model.

## V. Results

**Table 1:** Test accuracy on the size ranks and different simulated read lengths. Variable_len: Used variable length data augmentation, Fixed_len: Used fixed length data augmentation, MLP: Multilayer Perceptron, ResNet: Residual Network as an example of CNN, DC: direct sequence of characters, W2V: sequence of k-mers with Word2Vec embedding, SK: sequence of k-mers without Word2Vec.

|  | Training time<br>in mins | Prediction<br>time in secs | Species | Genus | Family | Order | Class | Phylum |
| --- | --- | --- | --- | --- | --- | --- | --- | --- |
| Variable_len_MLP_W2V | 172 | 18 | 0.7471 | 0.9626 | 0.9919 | 0.9947 | 0.9978 | 0.9984 |
| Variable_len_MLP_SK | 82 | 18 | **0.7412** | **0.9618** | **0.9911** | **0.9944** | **0.9976** | **0.9978** |
| Variable_len_MLP_DC | 28 | 18 | 0.1220 | 0.1356 | 0.1369 | 0.2032 | 0.3020 | 0.4002 |
| Variable_len_ResNet_W2V | 1660 | 174 | 0.7526 | 0.9641 | 0.9921 | 0.9953 | 0.9982 | 0.9987 |
| Fixed_len_MLP_SK | 26 | 18 | 0.0660 | 0.1356 | 0.0666 | 0.0776 | 0.3020 | 0.3412 |
| Fixed_len_ResNet_W2V | 1590 | 174 | 0.7426 | 0.9605 | 0.9920 | 0.9953 | 0.9980 | 0.9986 |

Recent CNN architectures such as ResNet and Inception achieved high accuracy at all levels till the genus level on the direct sequence of characters. On the other hand, they achieved ∼75% test accuracy on the species level for full-length HVR. ResNet was chosen as an example of CNN models to compare with MLP. This accuracy was the maximum result for all of our models. These models required at least 27 hrs of training to converge.

On the other hand, MLP achieved the maximum accuracy on W2V and SK with only ∼1.5 & 3 hrs respectively of training. Also, MLP models prediction time on Tesla K16 GPU was 10 times faster than ResNet. It’s worth mentioning that Word2Vec didn’t increase the accuracy nor the training time in all CNN networks. Despite Word2Vec allowed probable convergence to the maximum accuracy, whereas this didn’t happen in SK. Fixed length data augmentation didn’t allow MLP convergence but allowed for ResNet near the maximum accuracy.

**Table 2:** Best model test accuracies over different lengths and ranks. Full_HVR: The full length of the High variable region

|  | 25 | 50 | 75 | 100 | 125 | Full_HVR |
| --- | --- | --- | --- | --- | --- | --- |
| **Phylum** | 0.897 | 0.983 | 0.994 | 0.996 | 0.997 | 0.998 |
| **Class** | 0.862 | 0.976 | 0.991 | 0.995 | 0.997 | 0.998 |
| **Order** | 0.774 | 0.947 | 0.979 | 0.987 | 0.993 | 0.994 |
| **Family** | 0.726 | 0.922 | 0.966 | 0.981 | 0.988 | 0.991 |
| **Genus** | 0.588 | 0.818 | 0.894 | 0.930 | 0.950 | 0.962 |
| **Species** | 0.453 | 0.617 | 0.678 | 0.710 | 0.728 | 0.741 |

## VI. Discussion

Simple DL models such as MLP over SK allow 20X faster in training and 10X in prediction in comparison to CNN models. Moreover, the sequence of k-mers takes into account the k-mers position. We train each model to predict the 6 ranks in the same time because a multi-output model can learn better than a separate model for each output [48]. By comparing MLP models with SK and W2V, both converged near the maximum accuracy but SK was faster in training. On the other hand, Word2Vec prevented complex CNN models from over-fitting with a sequence of k-mers. Even this advantage didn’t achieve higher accuracy than DC but W2V with CNN might be useful in one-shot learning [49].

We model SILVA taxa, as it has more examples per class than SILVA nr, moreover, Greengenes latest update was on 2013. k-mer size of 6 was chosen empirically to provide unique k-mers but won’t generate a high number of k-mers that will increase the classifier complexity. In addition, Word2Vec corpus was saved locally to save memory during training the Word2Vec model. RNNs were the least explored architectures as they are vulnerable to over-fitting in case of the low number of samples per class.

Real-time data augmentation eliminate the need to generate all possible sub-sequences of a specific length, which will lead to increasing data size by at least 100 folds. Also, it allows generating new inputs for the model in each epoch to prevent the overfitting. Using HVR specific models decrease the search space during training, also it is usable in the application as any amplicon study should have known targeted HVR. ResNet model complexity may be the reason why fixed length data augmentation allowed ResNet convergence which is not the case in MLP.

## VII. Conclusion & Future Work

Most taxonomic classifiers depend on features rather than the direct sequence which can be modeled by DL. These features ignore their positions, resulting in low accuracy on short reads. Reducing the DL model complexity from CNN to MLP with global max pooling converged to species level classification with 74% test accuracy. MLP model decreased the training and prediction time by 20 and 10 folds respectively in comparison to CNN over a sequence of k-mers. Variable length data augmentation was the best data preprocessing for MLP besides its role in preventing overfitting and saving memory. Nevertheless we modeled only V2 HVR database, Future work will include modeling the rest HVRs databases. moreover, one-shot learning will be tested as many species class has very few samples.

**Figure 2:**
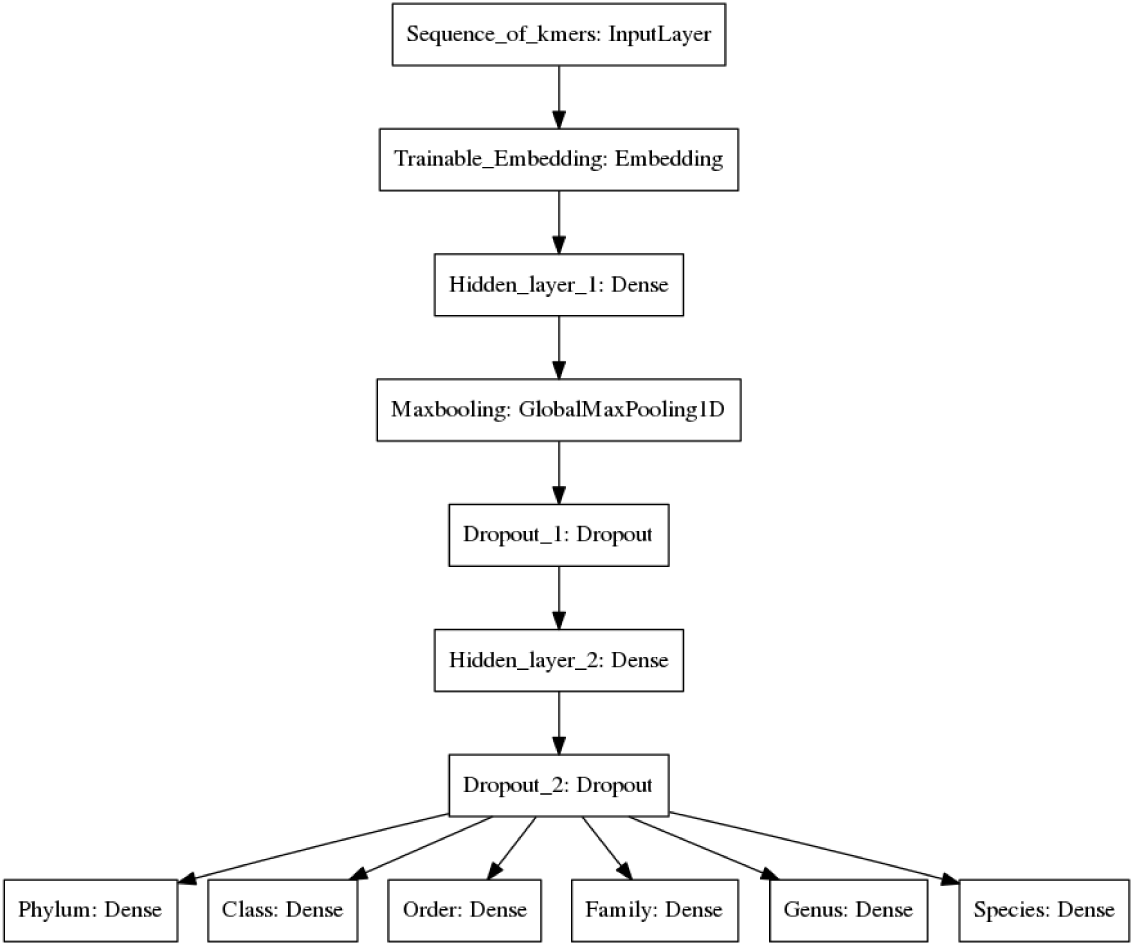
Best model architecture. The model begins with input layer to a sequence of k-mers, followed by trainable embedding layer, followed with 2 hidden fully connected layer with Global max pooling between them. Dropouts were added to prevent the overfitting. The model ends with 6 fully connected layers as outputs for each rank.

## Acknowledgment

Thanks for Karim Amer from the Ubiquitous & Visual Computing Group for his advices on architecture engineering. Most of the training was done on Google Colab GPUs.

